# D.sort: template based automatic spike sorting tool

**DOI:** 10.1101/423913

**Authors:** Madeny Belkhiri, Duda Kvitsiani

## Abstract

Understanding how populations of neurons represent and compute internal or external variables requires precise and objective metrics for tracing the individual spikes that belong to a given neuron. Despite recent progress in the development of accurate and fast spike sorting tools, the scarcity of ground truth data makes it difficult to settle on the best performing spike sorting algorithm. Besides, the use of different configurations of electrodes and ways to acquire signal (e.g. anesthetized, head fixed, freely behaving animal recordings, tetrode vs. silicone probes, etc.) makes it even harder to develop a universal spike sorting tool that will perform well without human intervention. Some of the prevalent problems in spike sorting are: units separating due to drift, clustering bursting cells, and dealing with nonstationarity in background noise. The last is particularly problematic in freely behaving animals where the noises from the electrophysiological activity of hundreds or thousands of neurons are intermixed with noise arising from movement artifacts. We address these problems by developing a new spike sorting tool that is based on a template matching algorithm. The spike waveform templates are used to perform normalized cross correlation (NCC) with an acquired signal for spike detection. The normalization addresses problems with drift, bursting, and nonstationarity of noise and provides normative scoring to compare different units in terms of cluster quality. Our spike sorting algorithm, D.sort, runs on the graphic processing unit (GPU) to accelerate computations. D.sort is a freely available software package (https://github.com/1804MB/Kvistiani-lab_Dsort).

## I. Introduction

Progress in systems neuroscience has been fueled by tools that record individual neurons in an intact brain. These tools rely largely on algorithms that classify extracellularly recorded electrophysiological signals into single units. When recorded on multiple channels each single unit shows unique spatial and temporal, spike waveforms assumed to originate from single neurons [1, 2].With the advancement of high density, high channel count electrophysiological recordings [**?**], there is a growing demand to develop accurate, fast, and automatic spike sorting tools [3–5]. However, inherent problems with extracellular recordings and a shortage of ground truth data [1,2] make it very difficult to establish a universal method for spike sorting, as evidenced by the variety of algorithms that have been developed recently [6]. The major problem with extracellular spike sorting tools, both for manual and automatic ones, is the lack of knowledge on the sources of background noise [7]. This noise is particularly problematic in the extracellular recordings of freely behaving animals where it may be comprised of the subthreshold activity of nearby neurons and the various electrical artifacts that originate due to animal movement [8].

Current spike sorting tools deal with the noise by discarding signals below a user-defined threshold. Manual algorithms (such as MClust from A.D.Reddish, Xclust from M.A. Wilson, and SpikeSort3D, an offline spike sorter from Neuralynx) and a majority of automatic spike sorting algorithms [6] differ mainly in the initial steps of signal processing. Manual clustering tools rely on the acquisition of filtered and threshold triggered signal. Automatic spike sorting tools process raw, unfiltered signal, but essentially apply the same filtering and thresholding procedures offline. Signal is filtered, in some cases denoised, and a threshold is applied to take snippets of data that may contain spikes. The signal that passes the threshold in both cases is further projected into a relevant feature space to separate single units [3, 5] or unique spatio-temporal spike waveform template are built to detect spikes [4]. The spikes that match the template by crossing a user-defined threshold are retained. Thus, in all of these approaches, user-defined thresholds determine whether spikes are retained or discarded. For neurons whose soma are close to the recorded electrodes, this might not pose any problems, but for neurons that are farther away, spikes may not be detected due to a change in background noise. Furthermore, even for spikes that always cross the threshold, a change in background noise level will result in spike amplitude variability that can make spike sorting difficult. This problem is aggravated in long-term recordings when the small but consistent displacement of electrodes in brain tissue (so-called ”drift”) causes the spike amplitude of some recorded neurons to increase or decrease [9]. Also, some neurons generate bursts of activity where spike amplitude decreases consistently on successive spikes within a burst.The bursting neurons are difficult to cluster, as they always appear as separate units in the feature space, and post processing steps are needed to merge them [10]. To minimize these problems, we decided to perform spike detection using NCC. Normalization prevents problems with changes in absolute spike amplitude and captures its unique spatiotemporal shape. More precisely, spikes are detected by running cross correlations between normalized spatiotemporal templates on the denoised, filtered, and normalized voltage trace. Normalization also provides us with metrics to see how close individual spikes are to their template and how far they are from other units. The algorithm is implemented on a GPU and can be run on a standard PC at a faster speed than the acquisition time on the 32-channel recorded system.

## II. Method

D.sort follows these steps: filtering and whitening the data in the preprocessing step, building the template waveforms to detect spike and merging similar templates. D.Sort is a spike sorter of the same class as Kilosort or Spikying circus since we use a template-matching algorithm. However, we point out that only the template initialization steps is similar to Kilosort [4].

### A. Preprocessing

In the preprocess steps, in order to remove the low frequency components and correlated noise, we filter and whiten the recorded data. For the filtering, we use a non-causal band-pass filter by applying a third order Butterworth filter at 300 Hz and 5, 000 Hz. For the whitening, we calculate the co-variance matrix ∑ from which the whitening matrix *W_h_* is deduced as follows:

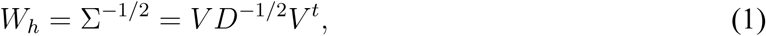

where *V* is a matrix containing the eigenvector of Σ and *D* is a diagonal matrix containing the eigen-value of Σ. Numerically, this calculation is achieved by adding a small constant *ϵ* = 10^−6^ in order to avoid a division by zero. Since the objective of whitening is to decorrelate the channels, we first detect and remove spikes by using a thresholding method:

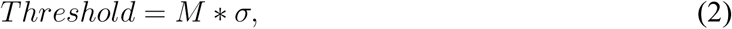

where *M* is an integer chosen by the user (equal to 4 by default) and *σ* is the standard deviation of the signal. After this step, we multiply the filtered data by *W_h_*, allowing the noise to be independent in time and space.

### B. Spike detection

In this step, the goal is to detect spikes and assign them to a cluster. To do this, we propose to detect spikes using NCC. The novelty introduced by the use of NCC is to focus on the waveform shape rather than relying on the spike amplitudes. Indeed, due to the various sources of noise (e.g. set-up, animal motion, thermal noise, etc.), the spike amplitude of a cell can vary, and normalization brings a solution to this problem. Therefore, using a normalized method allows us to avoid problems caused by base line shifts. Additionally, the NCC provides a score for each sampling point of how well the peak-aligned spikes fit to a given template. The NCC measure is

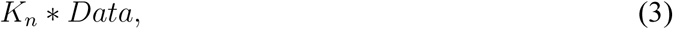

where *K_n_* is the template (*n* the number of templates) and *Data* is the filtered and whitened recorded voltage trace, both normalized. We point out that the template *K_n_* is defined on all recorded channels. This implies that we define a spike as a specific signal occurring on multiple channels. Therefore, each template is *N_channel_* × *N_sampling_*, with *N_channel_* being the number of channels and *N_Sampling_* being the number of sampling points.

#### Initialization

In order to apply the equation (3), we need initial templates to start detecting spikes. In the current version of D.sort, three alternatives initializations are possible. The first alternative uses artificial templates, as used by Kilosort [4]. The second alternative uses templates from a previous session. The goal of the latter is to track neurons across different recordings.

For the third alternatives, the templates are obtained from the data, where we use a simple thresholding method similar to the one presented at the whitening steps to detect spikes (cf equation (2). Then, those spikes are projected, channel by channel, onto a common base of artificial templates to get their *N* principal components. Next, a modified k-mean is applied using the *N* principal components with a user-defined number of *n* clusters. For more details about the cost function used in the modified k-means, we refer the reader to the paper on Kilosort [4]. Finally, the templates are deduced by averaging the waveform of their respective clusters.

#### Normalized cross correlation

As pointed out by Pachitariu et al.[4], computing the convolution with all the template components can be cumbersome and unnecessary. This is why, just as in Kilosort [4], we use a singular value decomposition (SVD) on the templates to extract their relevant features. Therefore, each template *K_n_* is defined by a sum of its *N* components, as shown below:

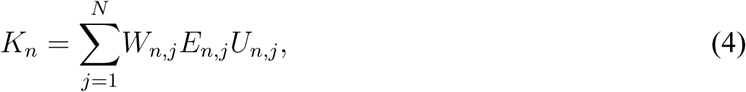

where *W* corresponds to the temporal components, *E* are the eigenvalues of the given components and *U* are the spatial components. For convenience and as it is in D.sort code, *W* will be *WE*, so *K* = *WU*.

#### Remarks

Since the matrix *U* contains the spatial information, it reports on which channel has the most relevant signal. *U* is an orthonormal unitary matrix, its value ranging from – 1 to 1. For example, in Figure 1, we show a template defined on four channels with the value of their first component in the spatial matrix *U*. As we can see, the strongest signal is on channel 15 with the value of 0.8, while the weakest is on channel 16 with *U* equaling 0.02. We clearly cannot rely on channel 16 to identify spikes, and therefore set a proper threshold criterion to disregard the irrelevant channels. Numerically, we evaluate the ratio of the U matrix on each channel with the maximum value of U. In the presented example, we would get in channel order the ratios: 0.3, 0.59,1,0.02. if we apply a threshold criterion of 0.3, we would set the U value of channel 16 to zero. Via this criterion, we define on which channel the spike is observed. The reason we set channels below a certain value to zero is to improve the computation by deleting noisy channels. As explained in the paper of Kadri et al. [11], the matrix *U* works like a mask that ignores irrelevant channels.

**Figure 1:**
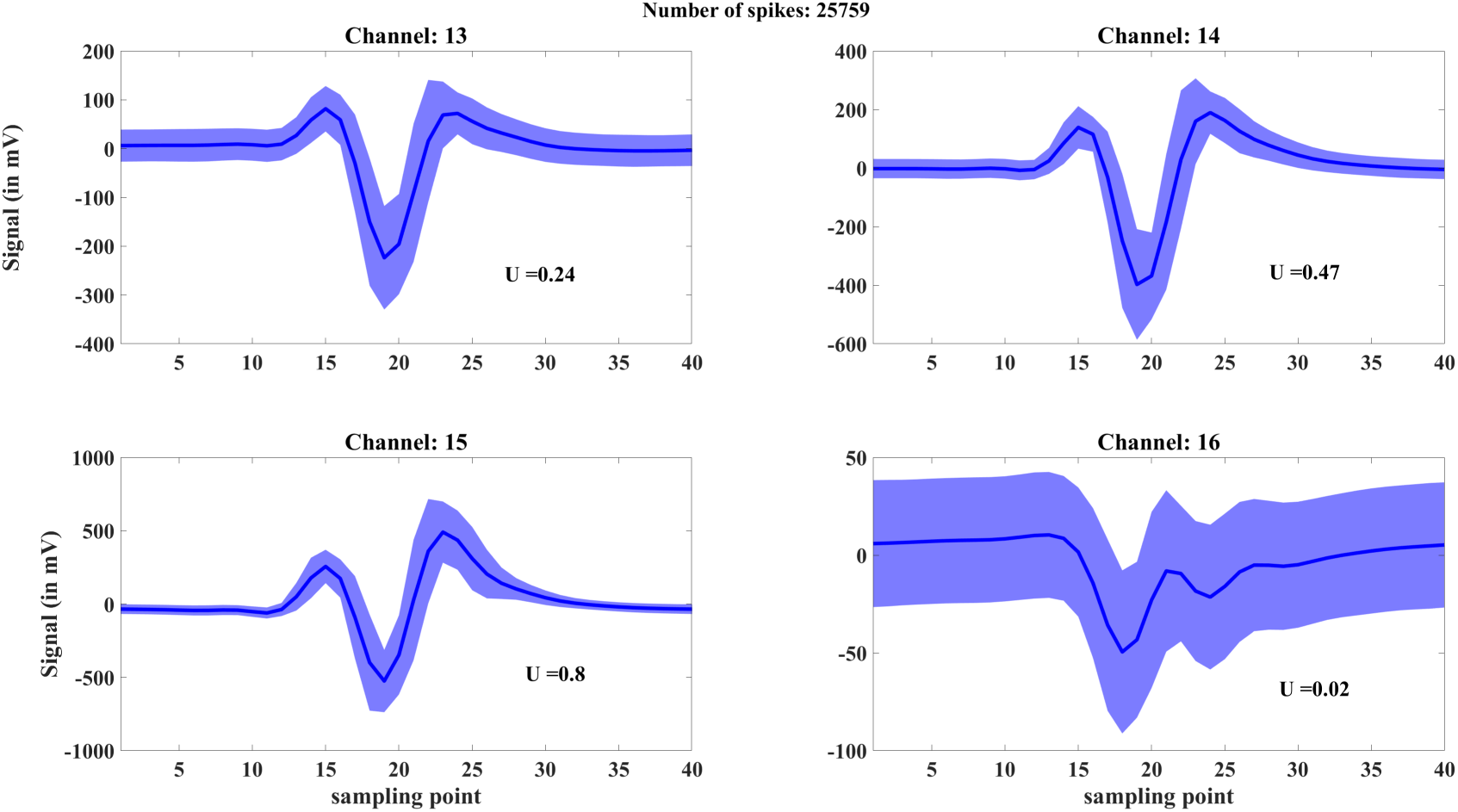
Mean waveform of cluster 7 from D.sort with its standard deviation for the channels of tetrode 3 with their respective first spatial component of the U matrix.

Using the SVD decomposition of equation (4), the NCC equation becomes

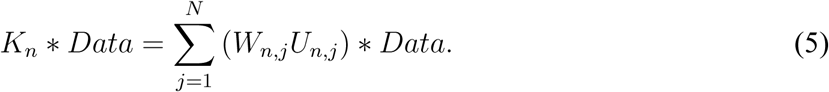

Since we have decomposed the templates into a sum of components, the NCC is also a sum of NCC for which the result is between –1 and 1. In the case of *N* = 3 components, the score of the NCC will be 3 at best. However, we do not find these results satisfactory, since each component of the templates captures a different amount of variance in the data set. Therefore, we weight each term of the NCC sum (cf equation (5)) by how much variance is captured by each component. In practice, for each template, we evaluate the captured variance for each component by calculating

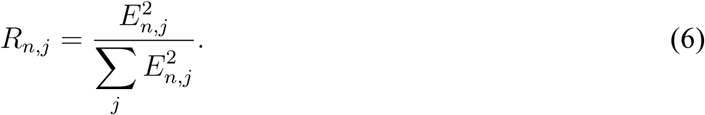

Then, we average the *R*_*n*,*j*_ over all templates to get the weights 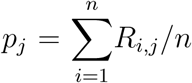 of each component. Finally, the computed formula for the NCC becomes

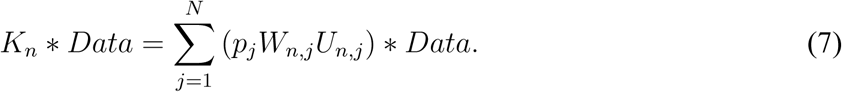

Our NCC version provides a number between –1 and 1 for each sampling point. Not only is this normalization used for detection, but it also gives a quantitative value of how well a spike matches a template without requiring any additional statistics.

In a computational perspective, the equation (7) is computed as follows:

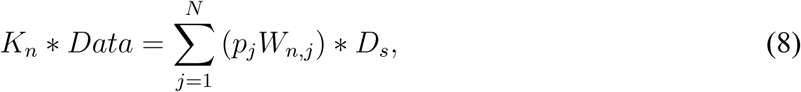

where *D_s_* is defined by

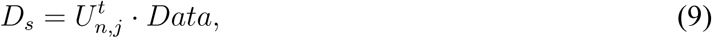

i.e., we first project the filtered data on the spatial components of *U* (where^*t*^ stands for the transpose). Then, we convolve *D_s_* with the temporal component *W*. We point out that the spatial projection is done on Matlab, while the convolution, which includes the normalization of *W* and *D_s_*, is done on CUDA. From equation (8), we get a matrix that we call *S* of dimension *n* × *M_s_*, where *n* is the number of templates and *M_s_* is the number of sampling point of the recording:

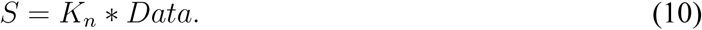

#### Timestamp extraction

In order to extract the timestamps, the matrix *S* is reduced into a *M* element vector labeled *Best*, by searching for the maximum score for each sampling point where

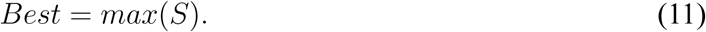

We apply a threshold to this vector to keep spikes that only match the templates reasonably well. The threshold value can be changed by the user (by default the value is set to 0.8). Then, we need to ensure that we do not count the same spikes twice. To do so, we write an algorithm that takes the first value of *Best* that is non-zero, and check that no over sampling point in the refractory period window is higher. If a score is higher in the window, then the algorithm moves to the next point and repeats the operation until no higher score is found.

#### Overlapping spikes

Here, we discuss the spatial and spatio-temporal overlap of spikes. The spatial overlap corresponds to spikes occurring at the same time (within 1ms) but being detected on different channels. The spatio-temporal overlap corresponds to spikes occurring at the same timestamps and being detected on the same channels but being emitted by different neurons. To circumvent these problems, it is has been proposed that the detected spikes be subtracted from the signal in order to uncover the other “hidden” spikes. The underlying assumption is that colliding spikes are simply a superposition of spikes. For spatio-temporal overlapping spikes, we note that this assumption is difficult to test due to the absence of ground truth data for in-vivo recordings. If spikes of different cells occur within the same millisecond but with some delay, we might detect them using such subtraction. However, if they occur with almost no delay, two problems arise: First, the resulting colliding spikes might be so deformed that the spikes would not pass the threshold criteria; and second, how should we subtract the template from these spikes? Should we peak align or base align?

For spatially overlapping spikes, the subtraction procedure is naturally efficient.

In our approach, we subtract the detected spikes from the signal, then we iterate in order to detect spikes that are spatially overlapping. The number of iterations is limited by the number of spikes detected by each iteration compared to the number of spikes already detected. By default, when an iteration detects less than *P*% (*P* being 2% by default) of the total number of spikes, the iteration is stopped.

##### Algorithm 1

Flowchart of spike detection

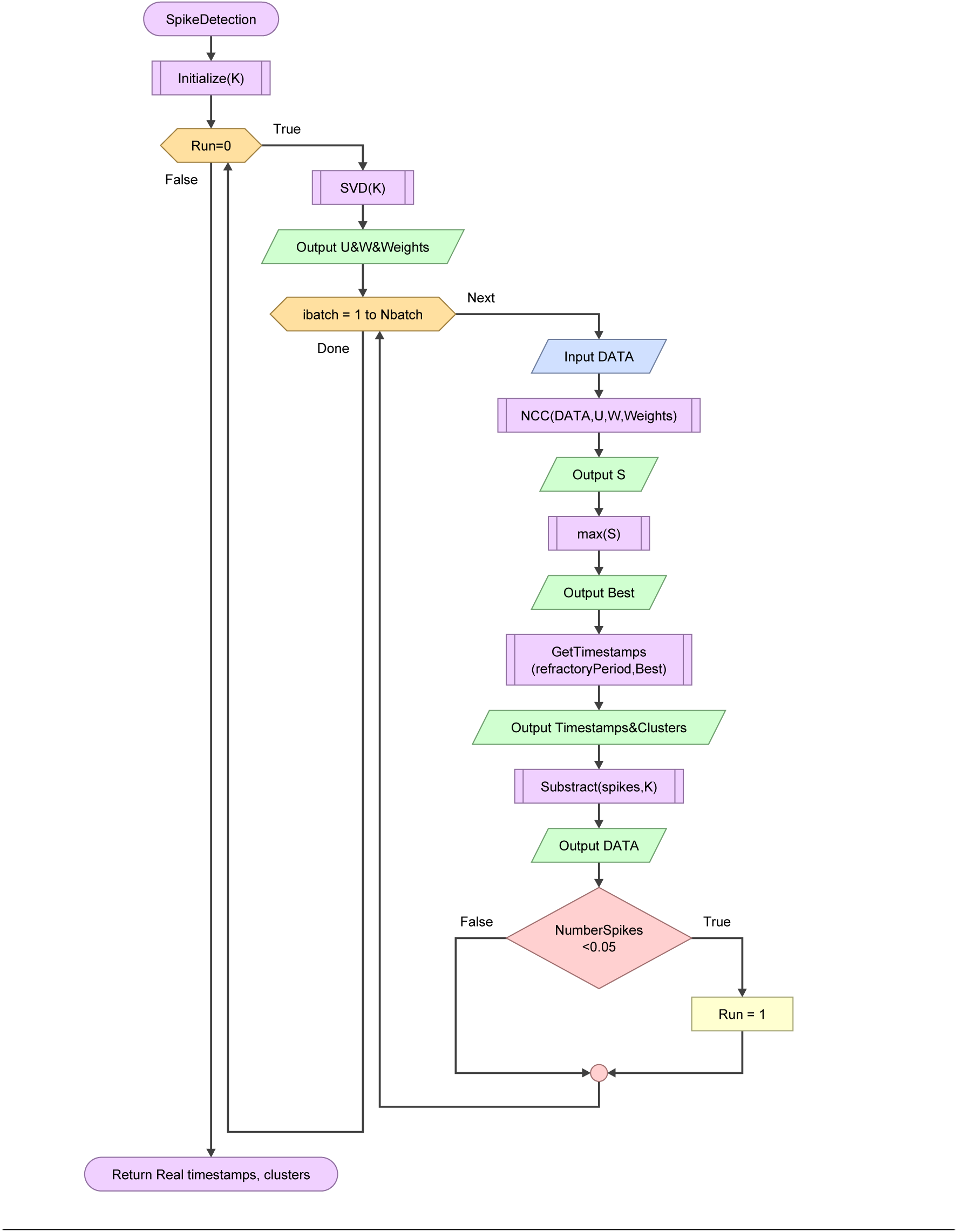

Finally, we summarize the above detection procedure in the following flowchart 1.

### C. Merging

To merge the clusters, we impose two conditions: Merged clusters should not violate the refractory period and they should have the same top channels in the U matrix (but not necessarily in the same order).

- Refractory period

In order to check that the refractory period is not violated, we construct an autocorrelogram with a bin size of 1 ms out of the merged clusters. Statistical significance with *p* < 0.01 of a dip in autocorrelogram around zero bin is evaluated by comparing it to the distribution of a temporally shuffled spike train over a window of 1 second. The surrogate distribution is generated by shuffling the timestamps of each spike 10,000 times.

- Top channels

The second condition is checked by using the first component of the *U* matrix. If the clusters exhibit the same top channel, they can be merged. If those two conditions are fulfilled, we use NCC on the mean waveform of the clusters and merge ones that get a score above 0.9.

## Acknowledgments

We thank Balázs Hangya, Jesper Hagelskjær, Szilard Sajgo, Junior Samuel Lopez Yepes, and all other members of the lab for providing feedback and the data. This work was supported by Lundbeck-fonden grant no.DANDRITE-R248-2016-2518.

